# Small extracellular vesicles reflect senescence progression in human bone marrow–derived mesenchymal stem cells during hollow fiber bioreactor culture

**DOI:** 10.64898/2025.12.31.697149

**Authors:** Ali Dinari, Armin M. Ebrahimi, Bartosz Leszczynski, Kamil Wawrowicz, Masoud Rezaei, Maciej Stotwinski, Zenon Rajfur, Ewa Stepien

## Abstract

Prolonged three-dimensional culture exposes stem cells to sustained microenvironmental and mechanical stresses that can promote aging- and senescence-associated phenotypic alterations. This study examined how long-term expansion of human bone marrow–derived mesenchymal stem cells (BMSCs) in a hollow fiber bioreactor (HFB) influences cellular senescence and the molecular composition of secreted small extracellular vesicles (sEVs). During extended HFB culture, BMSCs exhibited progressive morphological flattening and cytoskeletal disorganization, accompanied by increased senescence-associated β-galactosidase activity and immunophenotypic remodeling characterized by reduced fluorescence intensity and spatial redistribution of canonical MSC markers, consistent with a stress-adapted, early senescence–associated cellular state. In parallel, sEVs were collected longitudinally over 40 days and characterized by nanoparticle tracking analysis, immunoblotting, and quantitative proteomics. While vesicle size, marker expression, and yield remained stable throughout culture, proteomic profiling revealed pronounced, phase-dependent remodeling of sEV cargo, including coordinated alterations in oxidative stress–related processes, lysosomal and extracellular matrix–associated pathways, and relative depletion of cytoskeletal and translational components. Notably, these vesicular signatures closely mirrored senescence-associated changes observed at the cellular level. The strong correspondence between cellular phenotypes and sEV proteomic profiles establishes vesicle analysis as a convergent and noninvasive readout of BMSC aging, enabling sensitive monitoring of senescence progression while reducing reliance on parallel, labor-intensive cellular assays. Collectively, these findings indicate that prolonged HFB culture promotes a controlled, stress-associated senescence program in BMSCs and position sEV proteomic profiling as a robust approach for assessing stem cell aging dynamics during long-term three-dimensional bioreactor culture.

## Introduction

Cellular aging and senescence are fundamental biological processes that regulate tissue homeostasis, regenerative capacity, and intercellular communication. In mesenchymal stem cells (MSCs), senescence is characterized by progressive morphological enlargement, cytoskeletal remodeling, growth arrest, and altered expression of stemness-associated surface markers, ultimately impairing their functional potential and therapeutic utility (Boulestreau et al. 2020; Hayflick 1965). These age-associated phenotypic changes are increasingly recognized as dynamic processes influenced by cellular microenvironment, metabolic stress, and prolonged culture conditions. Extracellular vesicles (EVs) have emerged as key mediators of cell–cell communication and reflect the physiological state of their parental cells. In particular, small extracellular vesicles (sEVs) carry proteins, lipids, and nucleic acids that mirror cellular stress responses, aging-associated signaling pathways, and senescence-related remodeling (Théry et al. 2018). Accumulating evidence suggests that senescent or stress-adapted cells actively alter EV biogenesis and cargo composition, thereby propagating aging-related signals to surrounding cells and tissues(Théry et al. 2018). For clinical and translational applications, large-scale and standardized production of MSC-derived EVs is required(Gobin et al. 2021). Conventional two-dimensional (2D) culture systems are limited in scalability and reproducibility, prompting the development of three-dimensional (3D) culture platforms that better support long-term cell expansion. Hollow fiber bioreactors (HFBs) provide a controlled 3D microenvironment with enhanced nutrient exchange and waste removal, enabling sustained, high-density MSC culture with reduced manual intervention(Greuel et al. 2019; Watson et al. 2018; Lambrechts et al. 2016). HFB-based systems have been successfully applied for the production of recombinant proteins, viral vectors, and therapeutic nanovesicles, including MSC-derived EVs(Ala-Uotila et al. 1994; Yazaki et al. 2001; Jakl et al. 2023; Garcia et al. 2024). Despite these advantages, prolonged MSC culture, even under optimized conditions, can impose cumulative metabolic and mechanical stress that drives senescence-associated functional remodeling of both cells and their secreted vesicles(Boulestreau et al. 2020). While approaches such as genetic immortalization have been explored to overcome replicative senescence, MSCs inherently possess a finite lifespan, and culture-induced aging remains an unavoidable biological constraint (Hayflick 1965). However, how prolonged 3D culture shapes the senescence trajectory of MSCs and whether these changes are encoded within the molecular cargo of secreted sEVs remain incompletely understood. In this study, we investigated the effects of prolonged HFB cultivation on human bone marrow–derived MSCs (BMSCs), with a particular focus on senescence-associated cellular phenotypes and their relationship to longitudinal changes in sEV composition. BMSCs were expanded under conventional culture conditions and subsequently inoculated in HFBs for extended periods. Cellular aging was assessed using senescence-associated β-galactosidase staining, confocal microscopy, and flow cytometric immunophenotyping, while sEVs were characterized by nanoparticle tracking analysis, immunoblotting, and quantitative mass spectrometry. This integrated strategy enabled assessment of BMSC senescence at both cellular and vesicular levels, testing whether sEV molecular signatures provide a convergent readout of aging-associated cellular states under long-term 3D culture.

## Materials and methods

### Materials

Bone marrow derived mesenchymal stem cells, (Cat#:PB-CH-675-0511-44-1), mesenchymal stem cell medium Kit enhanced xeno-free (Cat#:PB-C-MH-675-0511-XF) with 25 ml growth supplement, and cryo-gold, sterile filtered, endotoxin tested, 100 mL (Cat#, PB-10003-01) were purchased from PELOBiotech (Germany). FBS (Cat#: FBS-H1-11A) purchased from Capricorn Scientific company (GmbH, Germany). Cell culture medium such as DMEM high glucose (Cat#: 61965026), xeno-free medium (Cat: PB-C-BH-675-0511-XF) were purchased from PELOBiotech and Gibco companies respectively. Accutase enzyme solution (Cat#: 637945), PBS (Cat#:10010023), and three layers cell culture Multi-Flask (#132867, DK-4000 Roskilde, Denmark) have been prepared accurately. Cartridge (Cat#: C2025D) purchased by Fiber Cell Systems (Maryland, USA). DAPI (4’,6-diamidino-2-phenylindole) (Cat# 28718-90-3) and Phalloidin–tetramethylrhodamine isothiocyanate (Phalloidin-TRITC) (Cat#: R415) were prepared from sigma. Cellular senescence assay kit (Cat#: KAA002) was prepared from merk (Poland).

### Methods

#### Conventional two-dimensional (2D) cell culture

in brief, a batch of cryopreserved BM-MSCs was thawed (within 1.30 minutes at 37 °C in a water bath), mixed with 5 mL of fresh xeno-free medium, centrifuged (8 minutes at 500g), and cultured in a T75 flask. Three passages of cell culture were performed to obtain an adequate number of cells. Next, the cells were subculture into 7 three-layer multi-flasks cell culture. The seeding density was 1.5 × 10⁶ cells per flask. At the proper confluency (90%) the cells were detached, centrifuged, resuspended in fresh xeno-free medium and transferred to intracapillary space (ECS) of HFBs.

#### Cell inoculation into HFBs

To inoculate the BMSCs into the HFBs, the harvested cells from conventional method were centrifuged, resuspended in fresh medium and transferred to the ECS of HFBs. In this regard, the proliferated BMSCs were treated with accutase enzyme solution, centrifuged at 500g for 8 minutes and resuspended in the fresh complete DMEM (10% FBS). The total volume of resuspended cells solution was 3ml and was pipetted gently to prevent cell clumping. Next, the cells were transferred to a sterilized syringe, attached to the side port of the cartridge, and injected into the ECS of HFBs. To ensure complete filling of the cartridge space with cells, the suspension was gently pushed back and forth five times (Supplementary Methods, S1, S2).

#### HFB Operation and Maintenance

HFB performance was monitored to assess the physiological status of inoculated BMSCs throughout culture. Key parameters included glucose consumption, cell viability, and sEV recovery (Supplementary Methods, S3). Culture conditions including medium supply, temperature, and gas exchange were checked daily to ensure stable operation. BMSCs were initially cultured in high-glucose DMEM supplemented with 10% FBS. At mid-run, the culture medium was transitioned to a 1:1 mixture of DMEM and xeno-free medium (total volume 175 mL) and circulated continuously through the HFB lumen using a Duet pump system. During pre-culture and the first three days, the flow rate was maintained at 20–25 mL/min and subsequently increased to 30 mL/min for the remainder of the run. The hollow-fiber membrane (20 kDa molecular weight cut-off) permitted bidirectional exchange of nutrients and metabolic waste while retaining EVs within the ECS. sEV-containing medium was harvested from the ECS four times per week, collecting 3-3.2 mL per sampling. Medium recovered from the ECS was also used for assessment of BMSC viability. Distinct collection procedures were applied for sEV isolation and viability analysis.

#### Glucose consumption

To assess cell metabolic status, monitoring glucose consumption serves as an effective indicator. Indeed, glucose levels in the reservoir bottle reflect cell metabolic activity. In this case, the medium was collected from reservoir bottle and assayed by glucometer. The glucometer displayed values in units of mg/dL.

#### Cell viability

In addition to monitoring glucose consumption, which can be used as an indirect indicator of cellular metabolism, the viability of reside cells in the ECS of HFBs can be assessed by subculturing the detached cells. To this end, the BMSCs were harvested from HFBs and resuspended in the fresh medium and were seeded to T25 flasks. Through this experiment, the potency of detached BMSCs was evaluated.

#### EVs isolation

The sEVs were isolated from 4 mL of culture medium collected from the ECS using a differential centrifugation and ultracentrifugation protocol. Briefly, the conditioned medium was centrifuged at 500 × g for 10 min to remove cells and large debris, followed by sequential centrifugation at 3,000 × g for 25 min and 7,000 × g for 30 min to eliminate residual cellular contaminants. The resulting supernatant was further clarified by centrifugation at 18,000 × g for 20 min. Finally, small EVs were pelleted by ultracentrifugation at 150,000 × g for 90 min. The sEV pellet was collected and used for downstream analyses (Supplementary Methods, S4).

#### Termination of HFBs operation

After a defined date, the HFBs operation system was terminated, and the cells were harvested and sub-cultured by conventional method. Through this stage, the effects of HFBs on cell morphology and viability were evaluated. The applied protocol for cell harvest is presented in Supplementary Methods (S5). In brief, the cartridge was treated with accutase enzyme, incubated at 37 °C for 1 hour and then flushed. Residual cells in cartridge’s ECS were detached and flushing with PBS. This step should be handled by swishing the applied enzyme rigorously between two connected syringes to detect the cells. The solvent was transferred to the 15ml conical tube and centrifuged with condition of 10 minutes at 500g. Next, the cells were enumerated and checked for their viability. Afterward, the cells were cultivated in T25 flask and incubated under optimum condition of incubator (37°C and 5% of Co2) and didn’t change the medium for three days. The process of cell proliferation and cell morphology were assessed correctly, and the results were recorded consequently.

### BMSC Characterization and Senescence Assessment

To evaluate the impact of HFB cultivation on BMSCs morphology three analytical techniques such as senescence-associated β-galactosidase (SA-β-Gal) assay, confocal microscopy, and flow cytometry were applied.

#### Cellular Senescence Assessment

Senescence was assessed using a commercially available SA-β-Gal staining kit following the manufacturer’s protocol. BMSCs were seeded in six-well plates at two densities: 5 × 10³ cells/well (≈20% confluency) and 2.5 × 10⁴ cells/well (≈80% confluency). Cells were fixed with 1 mL of 1× fixing solution for 15 minutes at room temperature, washed twice with PBS, and incubated with 2 mL of freshly prepared 1× SA-β-Gal detection solution at 37 °C (without CO₂) for 4 hours. After incubation, the solution was removed, cells were rinsed with PBS, and blue-stained cells were quantified under a phase-contrast microscope. Increased blue coloration indicated enhanced β-galactosidase activity and senescence.

#### Confocal Microscopy and Cell Staining

To visualize nuclear and cytoskeletal organization, DAPI and phalloidin staining were performed on fixed cells. DAPI stock solution (1 mg/mL) was diluted 1:1000 in PBS to a working concentration of 1µg/mL. Cells were incubated with DAPI for 10 minutes in the dark and rinsed twice with PBS. Permeabilization was achieved using 0.1% Triton X-100 for 10 minutes, followed by PBS washing and blocking with 3% BSA for 30–60 minutes at room temperature. Actin filaments were stained with Phalloidin–TRITC (100–200 nM) for 30–60 minutes in the dark. Cells were washed three times with PBS (5 minutes each) and imaged using a confocal laser scanning microscope. DAPI (excitation 358 nm; emission 461 nm) visualized nuclei, while Phalloidin–TRITC (excitation 540–555 nm; emission 570–580 nm) highlighted actin filaments.

#### Flow Cytometry

Immunophenotypic characterization of BMSCs was performed using a CellStream™ flow cytometer (Model: CS 073; Manufactured: October 2020; USA). BMSCs were detached from T75 flasks, washed with cold PBS, and resuspended in PBS containing 2% FBS. Approximately 5 × 10⁴ cells per sample were stained with 2 µL of each antibody in 100 µL PBS (2% FBS) for 30 minutes on ice in the dark. After two washes with PBS, the final cell pellets were resuspended in 200 µL PBS (2% FBS) for acquisition. The following antibodies were used for MSC characterization: CD105 (Cat# 2216090), CD90 (Cat# 2240590), CD73 (Cat# 2320020), and CD45 (Cat# 2120070) (Sony Biotechnology Inc.); and CD146 (Cat# 563619) and CD271 (Cat# 562122) (BD Biosciences).

##### Characterization of sEVs

To investigate the properties of BMSCs and their functional status during HFBs cultivation, sEVs were isolated and comprehensively characterized using nanoparticle tracking analysis (NTA), micro–BCA protein assay, western blotting, and high-resolution mass spectrometry (MS).

#### Nanoparticle Tracking Analysis (NTA)

The size distribution and concentration of sEVs were determined using a NanoSight NS500 system (Malvern Instruments, UK) equipped with a green laser and NTA software version 3.4 Build 3.4.4. The samples of sEV were diluted in sterile, filtered PBS to achieve optimal particle concentration. Analytical parameters were kept constant across all measurements (camera level 12, slider gain 245, detection threshold 2). The resulting data revealed overlapping size distributions typical of sEVs, with the predominant population exhibiting a hydrodynamic diameter consistent with exosomal vesicles.

#### Protein Quantification by Micro–BCA Assay

The total protein content of sEV preparations was quantified using the micro–BCA protein assay kit (Thermo Fisher Scientific, USA). Briefly, 5 µL of each sEV sample was mixed with 145 µL PBS, followed by 150 µL of micro–BCA working reagent. The mixture was incubated at 37 °C for 2 h, and absorbance was measured at 562 nm using a microplate reader. Protein concentrations were calculated against a bovine serum albumin (BSA) standard curve.

#### Western Blotting

Western blotting was performed to verify sEV identity and purity based on the presence of canonical vesicular markers. Antibodies against CD9 (Cat. #10626D), HSP70 (Cat. #sc-7298), and ARF6 (Cat. #sc-7971) were obtained from Santa Biotechnology (Japan). sEV lysates were prepared by mixing samples 1:1 with RIPA buffer and incubating on ice for 30 min with gentle agitation. Lysates were centrifuged at 13,500 rpm for 15 min at 4 °C, and the resulting supernatant was collected. Protein samples were mixed with 2× Laemmli buffer (1:1/4 v/v) containing 5% β-mercaptoethanol, heated at 100 °C for 7 min, and separated by SDS–PAGE (80 V stacking gel, 120 V resolving gel). Proteins were transferred onto PVDF membranes at 100 V for 1 h. Membranes were blocked in TBST containing 3% BSA for 1 h at room temperature, followed by overnight incubation at 4 °C with primary antibodies (1:1500 dilution in 1% BSA/TBST). After washing (3 × 5 min, TBST), membranes were incubated with rabbit anti-mouse HRP-conjugated secondary antibody (1:5000 dilution) for 1 h at room temperature. Chemiluminescent detection was performed using HRP substrate (Cat. No. WBKLS0100, USA), and signal acquisition was carried out using a qTouch Western Blot Imager (China). Expression of CD9, HSP70, and ARF6 confirmed the vesicular nature and integrity of the sEVs.

##### Proteomic analysis of sEVs by Mass Spectrometry

High-resolution proteomic profiling of sEVs was performed using a Vanquish Neo UHPLC system (Thermo Fisher Scientific, USA) coupled to an Orbitrap Astral mass spectrometer operating in data-independent acquisition (DIA) mode. Peptides were separated on a C18 analytical column under optimized gradient conditions. Raw DIA data were processed using DIA-NN v2.1.0 for spectral library generation, peptide identification, and label-free quantification. The human UniProtKB reference proteome (20,663 sequences; downloaded September 19, 2025) was used as the search database, with a 1% false discovery rate (FDR) applied at both peptide and protein levels. Data normalization, principal component analysis (PCA), and statistical evaluation were performed in R Studio (v2024.12.1 Build 563), and downstream bioinformatic analyses (hierarchical clustering, GO enrichment) were conducted using Perseus v2.1.5.0. Only proteins consistently identified across biological replicates were included in the final dataset, ensuring robust proteomic characterization of sEVs derived from HFB-cultured BMSCs.

#### Proteomic Data Processing

Quantitative proteomic profiling was performed on small extracellular vesicles (sEVs) isolated from BMSCs at three culture phases (Phase 1, Phase 2, Phase 3), with three biological replicates per phase. Protein abundance values were obtained from the proteomic software and exported together with statistical outputs, including fold change, p-values, q-values, and test statistics. Pairwise differential expression analyses were conducted using the built-in Student’s t-test framework of the proteomic platform.

#### Volcano plot generation

Volcano plots were generated for each pairwise comparison (Exo₂ vs Exo₁, Exo₃ vs Exo₁, Exo₃ vs Exo₂). Log₂ fold change (log₂FC) values were plotted on the x-axis and −log₁₀(p) values on the y-axis using Microsoft Excel. Proteins were categorized as upregulated (log₂FC > 1, p < 0.05), downregulated (log₂FC < −1, p < 0.05), or non-significant (p ≥ 0.05). Threshold lines marking log₂FC = ±1 and −log₁₀(p) = 1.3 were included to aid visualization. Differential expression categories were color-coded in all volcano plots.

#### Heatmap analysis

Heatmaps were produced to visualize expression patterns across the nine sEV samples using Morpheus (Broad Institute; https://software.broadinstitute.org/morpheus). Protein IDs and their normalized abundance values were extracted from the primary dataset, reorganized into a new Excel sheet, and subjected to row-wise z-score normalization. Log₂-transformed values were used for visualization. Hierarchical clustering of proteins and samples employed Euclidean distance and complete linkage. Color gradients represented relative abundance, enabling identification of coordinated phase-specific expression profiles.

#### Venn and UpSet Intersection Analyses

To examine shared and phase-specific differentially abundant proteins, proteins meeting the significance criteria (|log₂FC| ≥ 1 and p < 0.05) were compiled for each comparison. Venn diagrams were generated using InteractiVenn website (10.1186/s12859-015-0611-3) to illustrate simple overlaps, while UpSet plots were constructed using Intervene (Khan A, Mathelier A. Intervene, 2017) to resolve higher-order

#### STRING analysis

Protein–protein interaction networks were analyzed using STRING v12 (https://string-db.org/). Lists of differentially abundant proteins were uploaded, and interaction evidence incorporated experimental data, curated databases, co-expression, and text mining. A high-confidence interaction threshold (score = 0.7) was applied. Networks were visualized with nodes representing proteins and edges representing interaction confidence. STRING functional enrichment was used to identify significantly overrepresented GO terms, KEGG pathways, and molecular processes within each upregulated and downregulated set.

## Results and discussion

MSC senescence is a progressive process characterized by morphological remodeling, metabolic reprogramming, altered surface marker expression, and irreversible proliferative arrest (Li et al. 2017). Accumulating evidence indicates that senescence markedly compromises MSC therapeutic efficacy and the bioactivity of their secreted factors, including EVs (Neri & Borzì 2020). While genetic strategies such as immortalization have been explored to mitigate senescence, these approaches raise substantial biosafety and regulatory concerns and may alter fundamental MSC properties. HFBs represent a scalable and reproducible three-dimensional (3D) culture platform capable of supporting high-density MSC expansion under controlled perfusion conditions (Ren et al. 2024). By providing continuous nutrient exchange, a high surface-area-to-volume ratio, and reduced manual manipulation, HFBs improve process consistency compared with conventional two-dimensional (2D) culture (Hou et al. 2022). Moreover, 3D culture systems more closely approximate key aspects of the in vivo MSC niche(Bartosh & Ylostalo 2019). However, prolonged exposure to 3D mechanical constraints and sustained metabolic demand may impose cumulative stress, potentially accelerating senescence-associated phenotypic remodeling. This study investigated how extended HFB cultivation influences bone marrow–derived MSC (BMSC) physiology, senescence development, and extracellular vesicle output. We demonstrate that although HFBs support efficient initial cell expansion and metabolic stability, prolonged culture induces cytoskeletal reorganization, metabolic adaptation, and progressive loss of proliferative competence, consistent with a stress-associated senescent phenotype. These cellular changes are paralleled by pronounced remodeling of the secreted small EV (sEV) proteome, linking bioreactor-induced senescence to alterations in vesicle composition.

### Cell inoculation, glucose consumption, and viability in the HFB

BMSCs were efficiently inoculated into the HFBs at high density (∼6.5 × 10⁷ cells), achieving uniform adhesion and rapid establishment of a confluent cell layer along the fiber surfaces. Early-phase culture was characterized by high cell viability and stable proliferation, indicating effective adaptation to the perfused 3D microenvironment. These findings confirm that the HFBs supports robust initial MSC colonization and growth under controlled conditions. To assess metabolic stability during prolonged culture, glucose consumption was continuously monitored as a surrogate marker of cellular activity and viability. MSCs are adapted to low-oxygen niches and rely predominantly on glycolytic metabolism; thus, sustained glucose utilization reflects preservation of metabolic homeostasis(Moya et al. 2018). Throughout the 39-day culture period, glucose levels exhibited a steady-state consumption profile, indicating maintained metabolic activity despite increasing culture duration. Periodic medium exchange preserved nutrient balance, and a gradual transition to xeno-free conditions did not disrupt overall metabolic stability, underscoring the metabolic adaptability of MSCs within the HFB. Despite sustained glucose utilization, functional viability has progressively declined with extended culture duration (Figure 1, supplementary data,S3). Cells harvested at early time points readily reattached and proliferated following transfer to 2D culture, whereas late-stage harvested cells showed reduced attachment efficiency, diminished proliferative capacity, and increased cell loss. This dissociation between maintained bulk metabolic activity and declining regenerative competence suggests that prolonged HFB culture supports cellular survival but promotes cumulative mechanical and metabolic stress that compromises MSC functional integrity. Such stress-associated decline is consistent with early stages of senescence, in which metabolic activity persists despite loss of proliferative potential.

**Figure 1.**
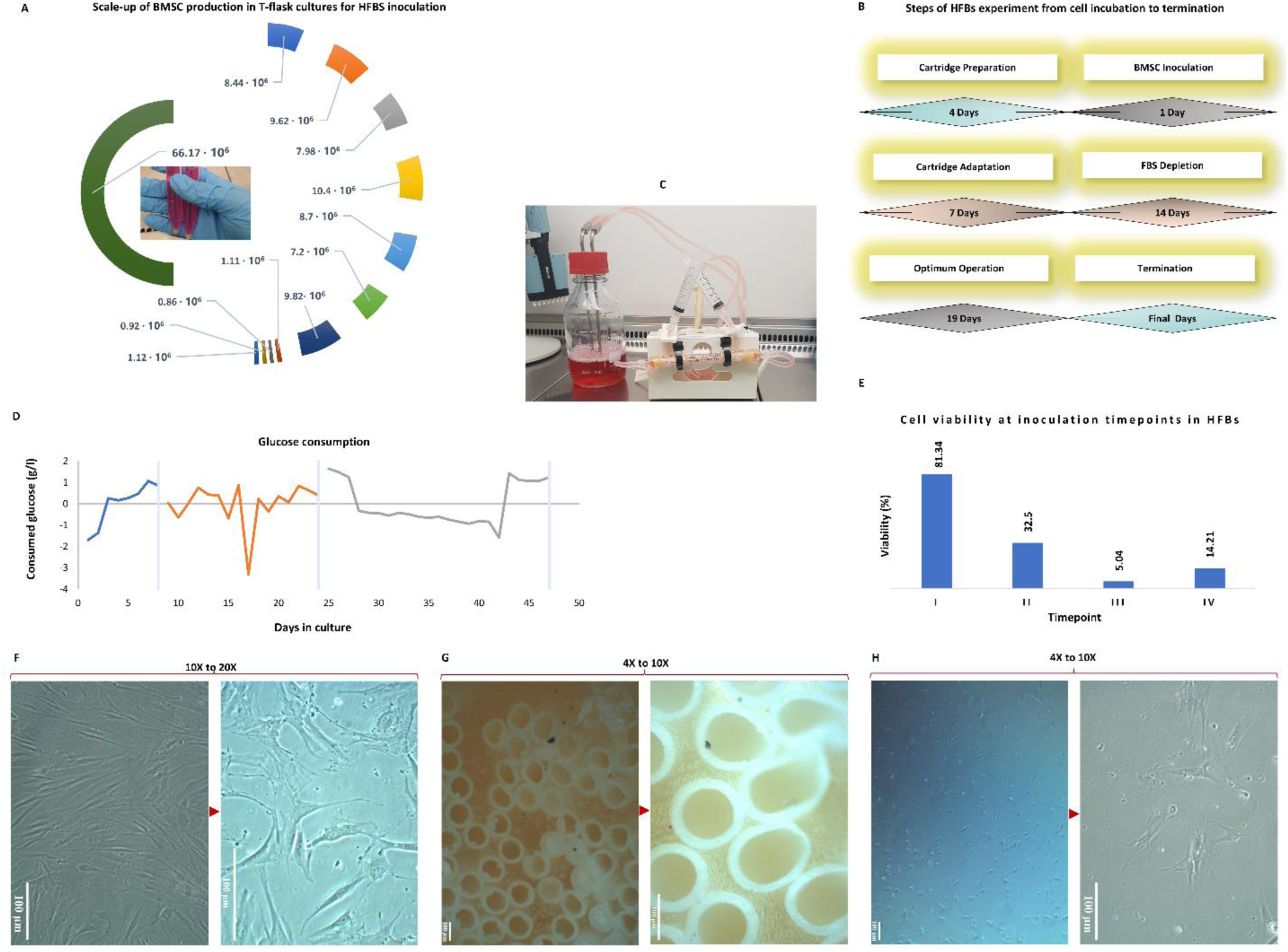
Scale-up, operation, and cellular responses of BMSCs during hollow fiber bioreactor (HFBS) culture. (A) Scale-up of human BMSC expansion in T-flasks prior to HFBS inoculation, illustrating cumulative cell yields obtained from different flask formats and passages used to achieve the required inoculation density. (B) Schematic overview of the HFBS experimental workflow, including cartridge preparation, cell inoculation, cartridge adaptation, stepwise fetal bovine serum (FBS) depletion, optimal operation phase, and system termination, with corresponding culture durations. (C) Representative image of the assembled HFBs setup during operation, showing the cartridge, tubing, and medium reservoir under continuous perfusion. (D) Glucose consumption profile over the course of HFBs culture, used as a surrogate marker of metabolic activity and viability. Shaded regions indicate distinct operational phases, including medium adaptation and extended culture. (E) Viability of BMSCs recovered from the ECS at defined inoculation/harvest time points, assessed after re-seeding under standard 2D conditions. (F–H) Representative phase-contrast microscopy images illustrating BMSC morphology under different culture conditions. (F) BMSCs maintained under conventional 2D culture display the characteristic elongated, spindle-shaped morphology (10×–20×). (G) BMSCs adhering to hollow fiber surfaces within the HFBs exhibit compact organization and altered spatial distribution (4×–10×). (H) BMSCs harvested from the HFBs and replated in 2D show flattened, irregular morphology and reduced cell density, consistent with senescence-associated remodeling (4×–10×). Scale bars are shown in each panel.

### Post-HFB harvest reveals senescence-associated morphological and functional remodeling

To further evaluate the impact of extended HFB culture on BMSCs phenotype, residual cells were recovered from the ECS and replated under standard 2D conditions (figure, F-H). Compared with the parental population, harvested cells exhibited marked morphological alterations, including loss of the characteristic elongated spindle shape, increased cell flattening, and irregular cell contours. These features are hallmarks of senescence-associated cytoskeletal remodeling and reduced mechanotransductive capacity. In parallel, harvested BMSCs displayed a pronounced reduction in proliferative activity and colony organization relative to control cells maintained exclusively in 2D culture. Decreased intercellular connectivity and disorganized growth patterns suggest impaired paracrine communication and altered secretory behavior following prolonged exposure to the HFB microenvironment. Given the central role of paracrine factors, particularly cytokines, growth factors, and EVs, in MSC-mediated tissue repair and immunomodulation, such changes are indicative of functional aging (Chimenti et al. 2010; Kizilay Mancini et al. 2017). Consistent with this interpretation, senescent MSCs are known to exhibit reduced paracrine potency and diminished therapeutic efficacy(Naji et al. 2019; Aggarwal & Pittenger 2005; Ferrucci & Fabbri 2018; Asumda & Chase 2011). Our observations indicate that prolonged HFB culture induces a phenotypic shift toward a senescence-associated state, characterized by preserved short-term survival but impaired regenerative and secretory function. These cellular alterations provide a mechanistic framework for understanding the progressive remodeling of sEV cargo observed during extended bioreactor operation.

### Characterization and senescence assessment of BMSCs cultured in HFBs

Senescence-associated β-galactosidase activity: To determine whether prolonged HFB culture induces cellular senescence, BMSCs harvested from the bioreactor were replated under standard conditions and assessed for senescence-associated β-galactosidase (SA-β-gal) activity (Figure 2-A). HFB-cultured BMSCs exhibited a marked increase in SA-β-gal activity, evidenced by intense cytoplasmic X-gal staining, whereas young, proliferative control BMSCs displayed negligible or absent β-galactosidase signal. SA-β-gal activity is a well-established hallmark of cellular senescence and reflects lysosomal expansion and altered metabolic homeostasis accompanying replicative and stress-induced aging. Consistent with prior studies, elevated SA-β-gal expression in BMSCs correlates with reduced proliferative capacity and progressive functional decline during cellular aging(Wu et al. 2014; Kim et al. 2012). The heterogeneous staining pattern observed among HFB-derived BMSCs indicates the coexistence of senescent and non-senescent subpopulations, supporting the concept that senescence develops progressively and asynchronously within expanding BMSC cultures. These findings demonstrate that extended HFB cultivation promotes the emergence of a senescence-associated phenotype rather than uniform terminal arrest.

**Figure 2:**
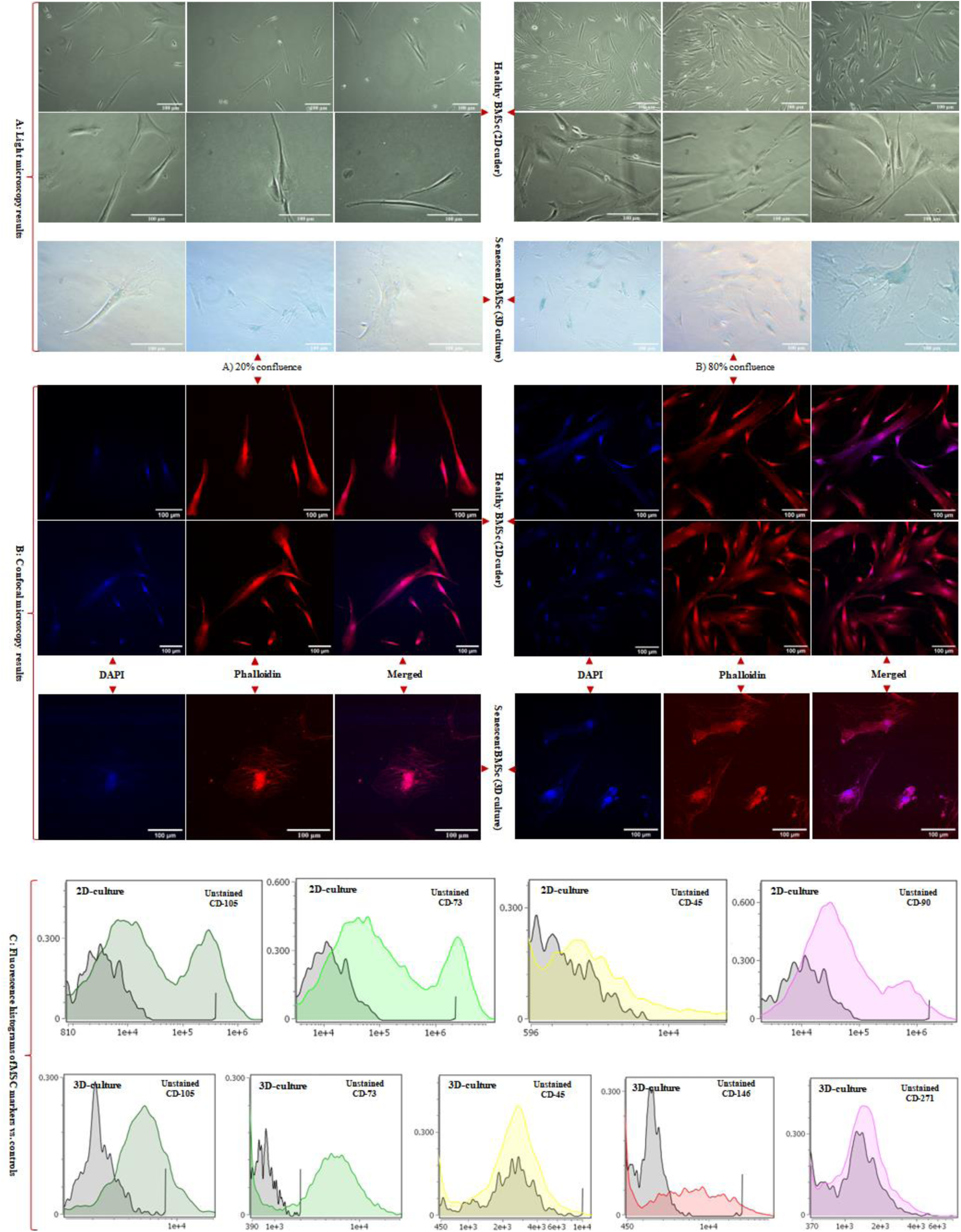
Representative light microscopy images show young BMSCs maintained under conventional 2D culture and senescent BMSCs harvested from the 3D hollow fiber bioreactor (HFB) system at low (20%) and high (80%) confluence. Senescence-associated β-galactosidase (SA-β-gal) activity was markedly increased in HFB-cultured BMSCs, visualized by characteristic blue-green X-gal staining at pH 6.0, compared with minimal staining in young controls. Confocal microscopy following DAPI (nuclei) and phalloidin (F-actin) staining revealed pronounced cytoskeletal disorganization, cell flattening, and nuclear enlargement in senescent cells relative to spindle-shaped, highly organized young BMSCs. All microscopy images were acquired using a 20× objective. Flow cytometric analysis compares immunophenotypic profiles of BMSCs cultured under 2D and HFB conditions. Histogram overlays depict fluorescence intensity distributions of antibody-stained samples relative to unstained controls. HFB culture was associated with reduced fluorescence intensity and altered distribution profiles of classical MSC markers (CD105, CD73), while CD146 expression was maintained or modestly increased and CD271 expression showed greater variability.

Cytoskeletal and nuclear remodeling revealed by confocal microscopy: To further characterize structural alterations associated with senescence, BMSCs were examined by confocal microscopy following DAPI and phalloidin staining to visualize nuclear morphology and F-actin organization, respectively (figure 2-B). Young BMSCs displayed the typical elongated, spindle-shaped morphology with well-organized, parallel actin filaments and uniformly shaped nuclei, reflecting preserved cytoskeletal tension and mechanotransductive signaling. In contrast, HFB-cultured BMSCs exhibited pronounced cytoskeletal remodeling, characterized by disrupted F-actin architecture, irregular filament accumulation, and loss of polarized organization. These cytoskeletal alterations were accompanied by nuclear enlargement, irregular nuclear contours, and reduced structural definition, indicative of compromised nuclear–cytoskeletal coupling and impaired cellular homeostasis. Such morphological features are characteristic of senescent MSCs and have been linked to altered mechanosensing, chromatin reorganization, and transcriptional dysregulation during cellular aging (Sun et al. 2024; Páez et al. 2020). Taken together, the combined increase in SA-β-gal activity and the observed cytoskeletal and nuclear abnormalities provide convergent evidence that prolonged HFB culture induces a stress-associated senescence program in BMSCs. Importantly, these structural hallmarks of cellular aging parallel the functional decline and phenotypic remodeling described earlier, reinforcing the conclusion that HFB conditions, while supporting prolonged survival and metabolic activity, impose cumulative mechanical and metabolic stress that drives senescence rather than sustained stemness.

Flow cytometry was employed to assess immunophenotypic changes in BMSCs maintained under conventional 2D culture and following prolonged cultivation in HFBs. Surface markers associated with MSC identity, purity, and functional heterogeneity (CD105, CD90, CD73, CD45, CD146, and CD271) were analyzed to evaluate culture-induced phenotypic modulation (Figure 2-C, Supplementary Table (S1, S2), Supplementary Figures (S1)). Under standard 2D conditions, BMSCs exhibited robust expressions of CD105, CD90, and CD73, with negligible CD45 positivity, fulfilling the minimal criteria for MSC identity defined by the International Society for Cellular Therapy (ISCT) (Dominici et al. 2006). Fluorescence intensity histograms showed pronounced rightward shifts relative to unstained controls, with occasional bimodal distributions, consistent with intrinsic heterogeneity within proliferative MSC populations. The absence of CD45 expression confirmed the lack of hematopoietic contamination (Bianco et al. 2013). In contrast, BMSCs cultured in HFBs exhibited marked immunophenotypic remodeling. While CD105 and CD73 expression remained detectable, both the proportion of positive cells and, more prominently, fluorescence intensity distributions were reduced compared to 2D controls. Histogram compression and attenuation of high-intensity tails suggested a contraction of highly expressing subpopulation rather than complete loss of MSC-associated markers. These changes are consistent with phenotypic convergence under prolonged 3D culture conditions, potentially reflecting adaptive responses to sustained perfusion, altered nutrient gradients, and mechanical constraints within the HFB microenvironment. Expression of CD146 was maintained or modestly increased following HFB culture, in line with its association with perivascular-like MSC subsets and sensitivity to microenvironmental cues. In contrast, CD271 expression was more variable and generally reduced, suggesting selective modulation of more primitive or stem-like MSC subpopulations during extended culture. Together, these marker-specific shifts indicate a redistribution of functional MSC subsets rather than uniform phenotypic loss. Taken together, these data demonstrate that HFB cultivation induces immunophenotypic remodeling of BMSCs characterized by attenuation and redistribution of classical MSC markers and altered representation of stemness-associated subpopulations. While such changes are compatible with a stress-adapted or early senescence–associated state, definitive attribution to irreversible senescence requires complementary functional analyses. Importantly, the phenotypic state of EV-producing cells is known to influence extracellular vesicle biogenesis and cargo composition (Théry et al. 2018), providing critical cellular context for the subsequent analysis of BMSC-derived small extracellular vesicles.

### Characterization of BMSC-derived sEVs produced in HFB culture

Approximately 70 mL of conditioned medium (3 mL per collection) was sequentially harvested from the ECS of the HFB system and subjected to comprehensive sEV characterization using nanoparticle tracking analysis (NTA), micro-BCA protein quantification, and western blotting. NTA measurements consistently revealed vesicle populations with a narrow size distribution predominantly below 200 nm, confirming enrichment of sEVs and excluding significant contamination by larger microvesicles or cellular debris. Quantitative comparison of particle concentration and total protein content demonstrated a strong positive correlation between vesicle number and protein yield, indicating proportional cargo loading across samples (figure 3, Supplementary Table (S3)). This relationship indicates that increases in sEV particle abundance are accompanied by corresponding increases in total vesicle-associated protein, reflecting coordinated regulation of vesicle release and bulk protein export. Importantly, this proportionality does not preclude substantial qualitative remodeling of sEV cargo, which is addressed in subsequent proteomic analyses. Notably, sEV output displayed a reproducible oscillatory pattern across the three defined operational phases of the HFB: (i) early cell adaptation (days 1–7), (ii) serum depletion (days 8–21), and (iii) transition to xeno-free conditions (days 22–40). Despite changes in medium composition and increasing culture duration, vesicle production remained sustained, with concentrations ranging from 3.9 × 10⁸ to 8.2 × 10⁹ particles/mL (mean ≈ 2.5 × 10⁹ particles/mL). Such rhythmic fluctuations are consistent with emerging evidence that EV secretion and cargo loading are influenced by stress-responsive and temporally regulated cellular pathways (Yeung et al. 2022; Bazié et al. 2021). In the context of prolonged HFB culture, these oscillations likely reflect adaptive secretory responses of BMSCs to cumulative metabolic, mechanical, and oxidative stress rather than simple growth-associated variation. Protein quantification by micro-BCA assay (Supplementary Figure, S2), revealed sEV protein concentrations ranging from 50 to 500 µg/mL, consistent with reported values for MSC-derived sEV preparations obtained under scalable culture conditions(Kowal et al. 2016; Théry et al. 2006). Importantly, maintained protein yields despite progressive cellular senescence suggest that secretory activity is preserved even as proliferative capacity declines, an observation aligned with the concept of a senescence-associated secretory phenotype (SASP), in which stressed or aging cells remain metabolically active while altering the qualitative nature of their secretome. Western blot analysis (Supplementary figures, S3) of representative sEV fractions confirmed the presence of established vesicular markers, including CD9, HSP70, and ARF6, detected at their expected molecular weights (∼25, ∼70, and ∼20 kDa, respectively). The concurrent detection of tetraspanin (CD9), cytosolic chaperone (HSP70), and membrane-associated trafficking protein (ARF6) verifies vesicle integrity, endosomal origin, and preservation of canonical sEV identity. These protein signatures are consistent with previously characterized BMSC-derived sEVs and validate the purity of the isolated vesicle populations (Muralidharan-Chari et al. 2009; Jeppesen et al. 2025; Théry et al. 2018). Taken together, these data demonstrate that the HFB system supports sustained and reproducible sEV production over extended culture periods, even as BMSCs undergo progressive senescence-associated phenotypic remodeling. Notably, the preservation of vesicle abundance and structural identity in the context of cellular aging indicates that HFB-expanded BMSCs retain secretory competence, while the qualitative evolution of sEV cargo addressed in subsequent proteomic analyses likely reflects adaptive, stress- and senescence-associated changes in the parental cell state. This integrated framework links prolonged bioreactor culture with MSC aging dynamics and phase-dependent remodeling of sEV content.

**Figure 3:**
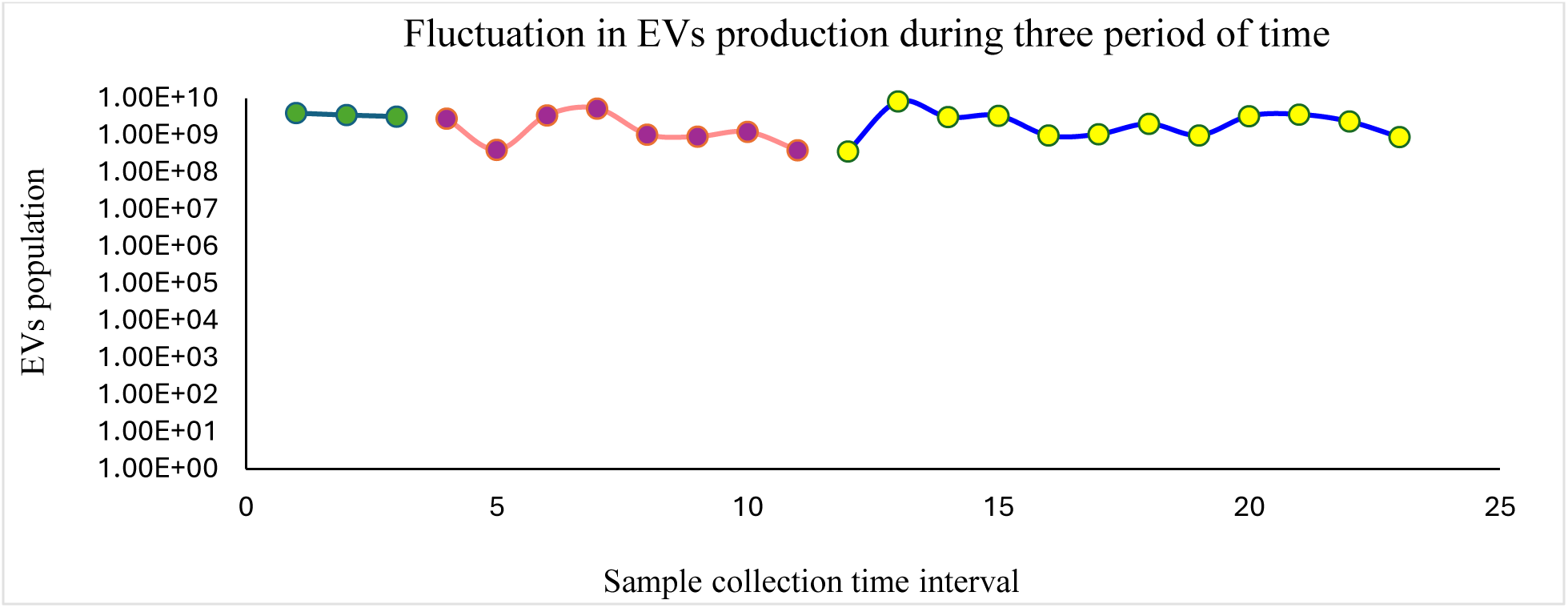
NTA analysis of EVs population (concentration) in samples volume.

### Proteomic analysis of sEVs by Mass Spectrometry

Primary analysis of quantitative proteomic data: Quantitative mass spectrometry revealed a progressive reduction in the number of proteins detected in sEVs across the three defined HFB culture phases, declining from Phase 1 (5,071 proteins) to Phase 2 (4,415 proteins) and further to Phase 3 (3,337 proteins) (Figure 4A). Despite this contraction in proteome breadth, differential expression analysis (Student’s *t*-test, *q* < 0.05) demonstrated extensive phase-dependent remodeling of sEV cargo. A total of 1,618 proteins were differentially regulated in Phase 3 versus Phase 1, compared with 579 in Phase 2 versus Phase 1 and 550 in Phase 3 versus Phase 2 (Figure 4B), indicating that the most pronounced proteomic divergence occurred between early and late culture stages. The overall reduction in detectable proteins by Phase 3 is therefore more consistent with cumulative culture-associated stress and the emergence of senescence-associated phenotypes in HFB-expanded BMSCs than with a decline in vesicle production itself. Cellular senescence is known to alter intracellular trafficking, translational capacity, and endolysosomal dynamics, resulting in selective restriction and reprogramming of EV cargo composition (Lim et al. 2021; Gobin et al. 2021). To further resolve the structure and functional implications of these proteomic changes, complementary analyses including volcano plots, hierarchical clustering heatmaps, overlap analyses, and functional enrichment using STRING were performed. The results of these analyses are detailed below.

**Figure 4.**
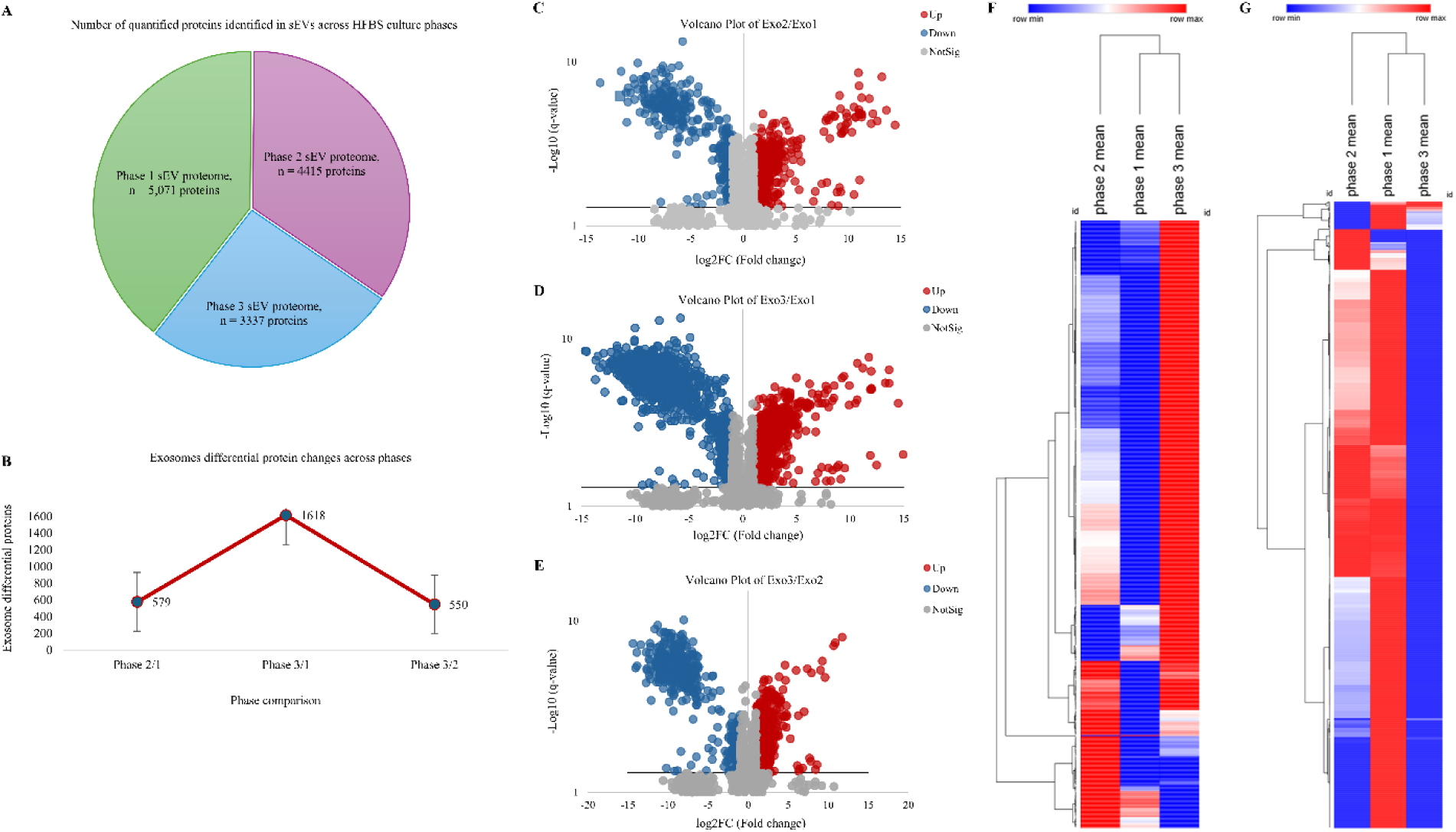
Differential exosome protein expression across experimental phases. Line plot showing the number of exosomal proteins with significant differential abundance between sequential culture phases (Phase 2/1, Phase 3/1, and Phase 3/2). Error bars represent variability across replicates. The largest proteomic shift occurred between Phase 3 and Phase 1 (1618 proteins), followed by Phase 2/1 (579 proteins) and Phase 3/2 (550 proteins), indicating dynamic remodeling of the exosome proteome across the culture timeline. Differential protein expression across experimental phases and functional network. (A–C) Volcano plots showing phase-wise differential protein abundance (A: Exo2 vs. Exo1; B: Exo3 vs. Exo1; C: Exo3 vs. Exo2). Red and blue points indicate significantly upregulated and downregulated proteins, respectively; grey points denote non-significant changes (|log₂FC| and adjusted p-value thresholds applied). (D–E) Heatmaps of significantly regulated proteins across phases. Panel (D) highlights proteins progressively upregulated toward later phases, whereas (E) shows proteins with inverse or phase-specific downregulation. Rows represent proteins; columns show mean phase expression; colors reflect row-wise z-scores (red: higher; blue: lower). (F–G) STRING protein–protein interaction networks for significantly upregulated (F) and downregulated (G) proteins. Dense subnetworks reveal enriched functional modules and coordinated regulation across phases, indicating biologically meaningful interaction hubs.

**Figure 5:**
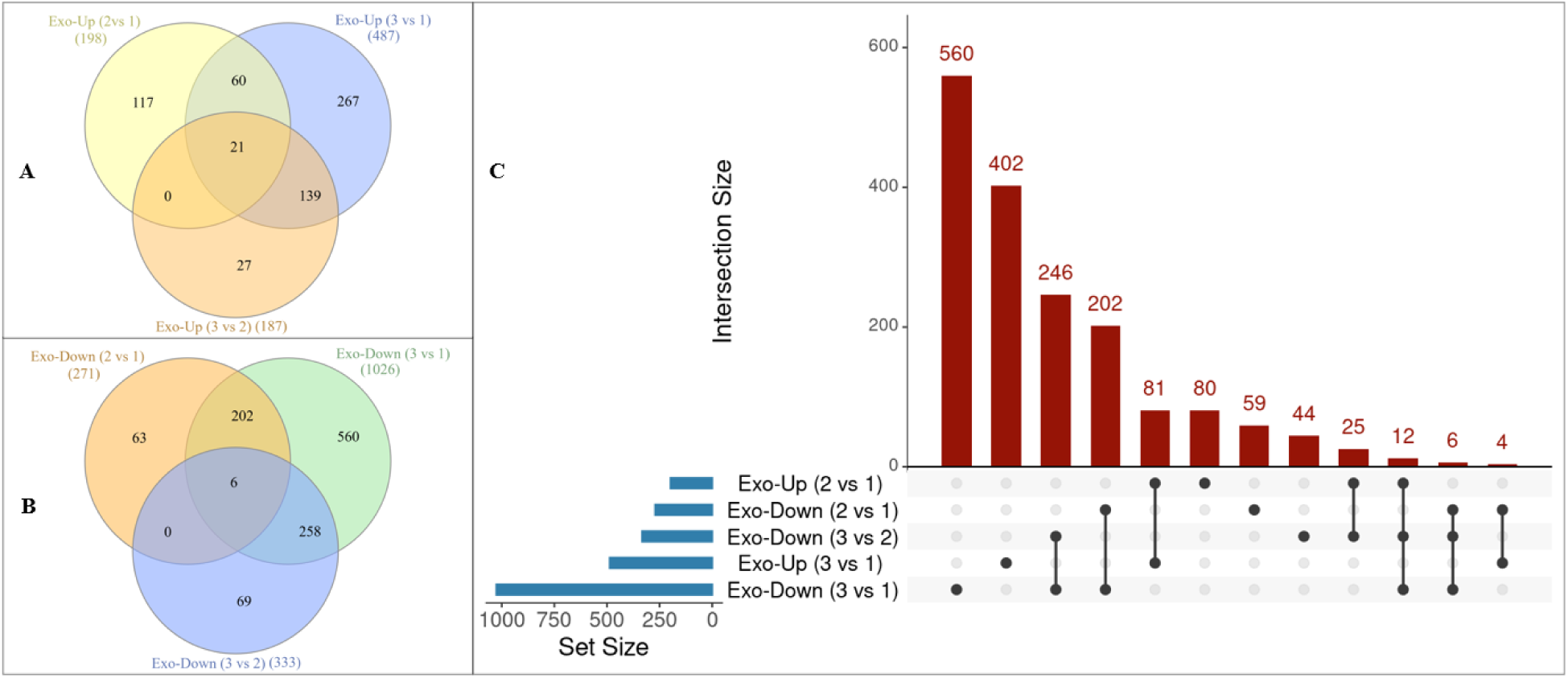
Venn and UpSet analyses of differentially expressed exosomal proteins across culture phases. (A–B) Venn diagrams showing overlaps among pairwise phase comparisons for upregulated (A) and downregulated (B) exosomal proteins. The three comparisons share 21 commonly upregulated and six commonly downregulated proteins. (C) UpSet plot summarizing set and intersection sizes. The largest unique intersections were Exo-Down (3 vs 1) (560 proteins) and Exo-Up (3 vs 1) (402 proteins), with additional major intersections indicated by bar labels. Together, these distributions highlight extensive phase-dependent and shared remodeling of exosomal cargo.

### Volcano–Heatmap–STRING analysis reveals phase-dependent remodeling of BMSC-derived sEV cargo

Comparative proteomic profiling across HFB culture phases demonstrated marked, coordinated remodeling of sEV protein cargo (Figure 4C–G). Volcano plot analyses revealed numerous significantly regulated proteins in each pairwise comparison, with the greatest divergence observed between Phase 1 and Phase 3 samples, consistent with progressive cellular adaptation to prolonged 3D culture. Hierarchical clustering of the union-regulated proteins resolved the data into two principal expression modules: one progressively enriched toward Phase 3 and another progressively depleted, indicating structured activation and suppression of distinct biological programs rather than stochastic proteomic drift(Gatto & Lilley 2012; Aebersold & Mann 2016). STRING network analysis (confidence score = 0.7, FDR = 0.05, MCL inflation = 1.8) identified 71 upregulated and 148 downregulated functional clusters (≥5 proteins per cluster). Proteins enriched in late-phase sEVs were predominantly associated with extracellular matrix organization, adhesion signaling, vesicle biogenesis, and endolysosomal pathways, alongside immune and stress-responsive signaling modules, including RHO–ROCK, mTOR-associated signaling, and ER protein processing. Evidence of metabolic reprogramming was also apparent, with relative enrichment of lipid-handling and glycolytic pathways, whereas components of oxidative phosphorylation and several amino acid metabolic pathways were comparatively underrepresented. In contrast, proteins progressively depleted from sEVs over time were enriched for nuclear and biosynthetic processes, including RNA splicing, ribosome biogenesis, RNA polymerase II–mediated transcription, nucleosome organization, and core metabolic functions such as the TCA cycle and branched-chain amino acid degradation. The selective loss of these biosynthetic and transcriptional regulators is consistent with reduced proliferative capacity and translational output characteristic of aging and pre-senescent MSCs. These coordinated shifts indicate that late-phase sEVs are not globally enriched but instead selectively repackaged, favoring stress-adaptive, ECM-associated, immune-related, and vesicle-trafficking proteins, while progressively excluding proteins linked to growth, biosynthesis, and metabolic homeostasis. This pattern aligns with senescence-associated reprogramming of MSC secretory activity, including mTOR-linked metabolic remodeling, endolysosomal and exosome-biogenesis adaptation, altered mechanotransduction and adhesion signaling and the emergence of innate and inflammatory signaling components reminiscent of SASP-associated EV cargo (López-Otín et al. 2013; Larsen et al. 2006; Herranz et al. 2015; Jeppesen et al. 2025; Salminen et al. 2012; Freund et al. 2012; Stewart et al. 2020). Additional enrichment of pathways related to TGF-β signaling, DNA damage response, and stress-associated cell death further supports a transition toward a senescence-influenced secretory state that dynamically reshapes sEV composition during prolonged HFB culture (Gaur et al. 2017; Ahmad et al. 2024).

### Phase-dependent sEV proteome remodeling during prolonged HFB culture reflects progressive MSC aging

Venn and UpSet analyses were employed to quantify shared and phase-specific differentially expressed proteins across the three HFB culture phases (Figure 3). Both analyses revealed a strong predominance of phase-specific changes, with the largest unique intersection observed for downregulated proteins in Phase 3 versus Phase 1 (Exo-Down 3 vs 1; 560 proteins), followed by uniquely upregulated proteins in the same comparison (Exo-Up 3 vs 1; 402 proteins). In contrast, comparisons involving Phase 2 showed smaller unique sets, consistent with a gradual rather than abrupt remodeling of the sEV proteome during prolonged culture. Only a limited core of proteins was conserved across all phase comparisons: 21 proteins were consistently upregulated and 6 proteins consistently downregulated across Exo-2 vs 1, Exo-3 vs 1, and Exo-3 vs 2. The small size of these shared intersections, relative to the extensive phase-specific sets, indicates that sEV cargo composition is highly sensitive to cumulative culture duration and microenvironmental transitions, rather than reflecting a static MSC secretory phenotype. This pattern is consistent with previous reports demonstrating that cellular stress and senescence drive selective, context-dependent remodeling of exosomal cargo rather than uniform shifts in protein abundance(Coppé et al. 2008; Mathieu et al. 2021). Mechanistically, the dominance of Phase 3 specific differentials suggests progressive rewiring of endosomal sorting and cargo-selection pathways as BMSCs transition toward a stress-adapted, senescence-associated state. Such selective loading is increasingly recognized as a hallmark of senescent EV biogenesis, whereby vesicles act as reporters of intracellular remodeling rather than passive byproducts of secretion (Basisty et al. 2020). Functional annotation of the 21 consistently upregulated proteins revealed enrichment for extracellular matrix (ECM) organization and structural remodeling components, including fibulins (FBLN1, FBLN4), tenascin (TNC), collagens (COL4A2, COL6A1, COL6A3), microfibril- and adhesion-associated proteins (AEBP1, SNED1, SVEP1), and matrix-modifying enzymes (OMD, CPXM2). This ECM-dominant signature indicates reinforcement of cell–matrix interactions and structural stabilization, processes closely associated with senescence-associated secretory phenotypes (SASP) and reduced regenerative plasticity. The presence of LMNB1 within sEVs further suggests nuclear-envelope stress and chromatin reorganization, features characteristic of early-to-intermediate senescence states(Freund et al. 2012). Concurrent enrichment of NUCB1 and DNAJC3 supports increased export of ER-stress and proteostasis-related signals via sEVs (Tsukumo et al. 2007). In contrast, the 6 consistently downregulated proteins were enriched for membrane-associated trafficking, transport, and signaling components, including CLDN11, LSAMP, RAI3/GPRC5A, FLOT1, and SLC38A1. Reduced abundance of FLOT1, a lipid-raft scaffolding protein implicated in non-clathrin endocytosis and exosome biogenesis(Fan et al. 2019), together with decreased SLC38A1, a glutamine transporter(Zhang et al. 2024), suggests attenuation of vesicle trafficking efficiency and altered metabolic support. Downregulation of adhesion- and membrane-organizing proteins further indicates remodeling of sEV membrane composition as senescence progresses (Liu et al. 2016). Notably, the opposing trajectories of ECM-associated proteins (e.g., fibulins, collagens) and membrane/trafficking regulators (e.g., flotillins) highlight a shift from an active, dynamic vesicle-sorting phenotype toward enhanced matrix stabilization and structural signaling. This transition aligns with established models of senescence-associated cytoskeletal reinforcement, extracellular matrix remodeling, and functional decline in long-term MSC cultures (Wiley & Campisi 2021).

### Transient phase-dependent exosomal signatures define multistage senescence adaptation

Venn-intersection analysis revealed discrete subsets of transiently regulated sEV proteins exhibiting directionally opposing expression across culture phases rather than monotonic change, indicating stage-specific remodeling of exosomal cargo during prolonged HFB culture. Heatmap analysis (Figure 6) confirmed pronounced direction-switching behavior, with proteins transiently induced during intermediate phases and subsequently suppressed at later stages, consistent with adaptive but ultimately constrained cellular responses during senescence progression(Coppé et al. 2008; Mathieu et al. 2021). One dominant protein module comprised factors linked to cell division, RNA metabolism, and adhesion-dependent mechanotransduction, including HMGB1, NOP56, TCOF1, U5-20k (U520), SRSF1, RBM25, GALT, STX16, CERT, LRRC15, RBP1, BKRB1, CLDN11, LSAMP, RAI3, SLC38A1, and FLOT1. Proteins involved in ribosome biogenesis and RNA splicing (NOP56, TCOF1, SRSF1, RBM25, U520) are essential for proliferative capacity and are known to decline during senescence-associated cell-cycle arrest and reduced biosynthetic output(Liang et al. 2023; López-Otín et al. 2013). Concurrent downregulation of adhesion- and membrane-associated proteins governing lipid-raft organization, junctional stability, and mechanotransductive signaling (e.g., FLOT1, CLDN11, LSAMP, STX16, CERT) indicates early disruption of cytoskeletal tension sensing and cell–matrix communication, hallmark features of pre-senescent mechanobiological stress responses (Han et al. 2025; Larsen et al. 2006). A second transient module displayed oscillatory regulation, characterized by suppression in Phase 1, induction in Phase 2, and renewed downregulation in Phase 3. This group included proteins involved in metabolic homeostasis, redox buffering, membrane trafficking, and paracrine signaling, such as MAT2A (METK1), ALDH1L1, PGAM4, KYNU, GSTA1/2, XDH, ALDH8A1, DPYS, SEC20, PLA1A, AREG, NOG, PTHR1, VEGFR3 (FLT4), and ANGPT1. These pathways regulate glycolytic flux, one-carbon metabolism, antioxidant defense, vesicular transport, and growth factor signaling, all of which are dynamically modulated during early senescence as cells attempt to buffer mitochondrial and cytosolic stress(Koh et al. 2020). Their transient induction during the intermediate phase suggests an adaptive stress-mitigation program that becomes progressively disengaged as cells transition toward a stabilized senescence-associated state. A smaller subset exhibited delayed re-expression at late culture stages, including RTN1, SHH, ANGPT1, and CA13, implicating compensatory ER–Golgi remodeling, morphogen signaling, vascular/paracrine regulation, and pH-linked metabolic adjustment during advanced senescence(Ahmad et al. 2024; Freund et al. 2012). Collectively, these data demonstrate that BMSC senescence under prolonged HFB culture proceeds through multistage, adaptive remodeling of exosomal pathways rather than a linear degenerative process. Early loss of proliferative and adhesion programs is followed by transient metabolic and paracrine compensation, culminating in stabilization of a senescence-associated secretory phenotype. The presence of reversible and phase-specific sEV signatures underscores the plasticity of MSC aging trajectories and establishes sEV proteomics as a sensitive, integrative readout of senescence stage, mechanotransductive stress adaptation, and functional decline in bioreactor-expanded BMSCs (Basisty et al. 2020; Salminen et al. 2012). Senescence progression under prolonged HFB culture is therefore more accurately described as a continuous, quantitatively regulated biological process rather than a discrete phenotypic transition. Whereas conventional cellular assays primarily capture integrated or downstream manifestations of stress adaptation, longitudinal sEV proteomic profiling resolves coordinated, multiaxial remodeling across metabolic, cytoskeletal, redox, and extracellular matrix–associated protein networks. The concerted regulation of functionally interconnected protein modules indicates that senescence reflects a system-level cellular reprogramming event. In this context, sEVs act as integrative molecular reporters of intracellular aging dynamics, providing higher-resolution insight into senescence trajectories than endpoint cellular measurements alone.

**Figure 6.**
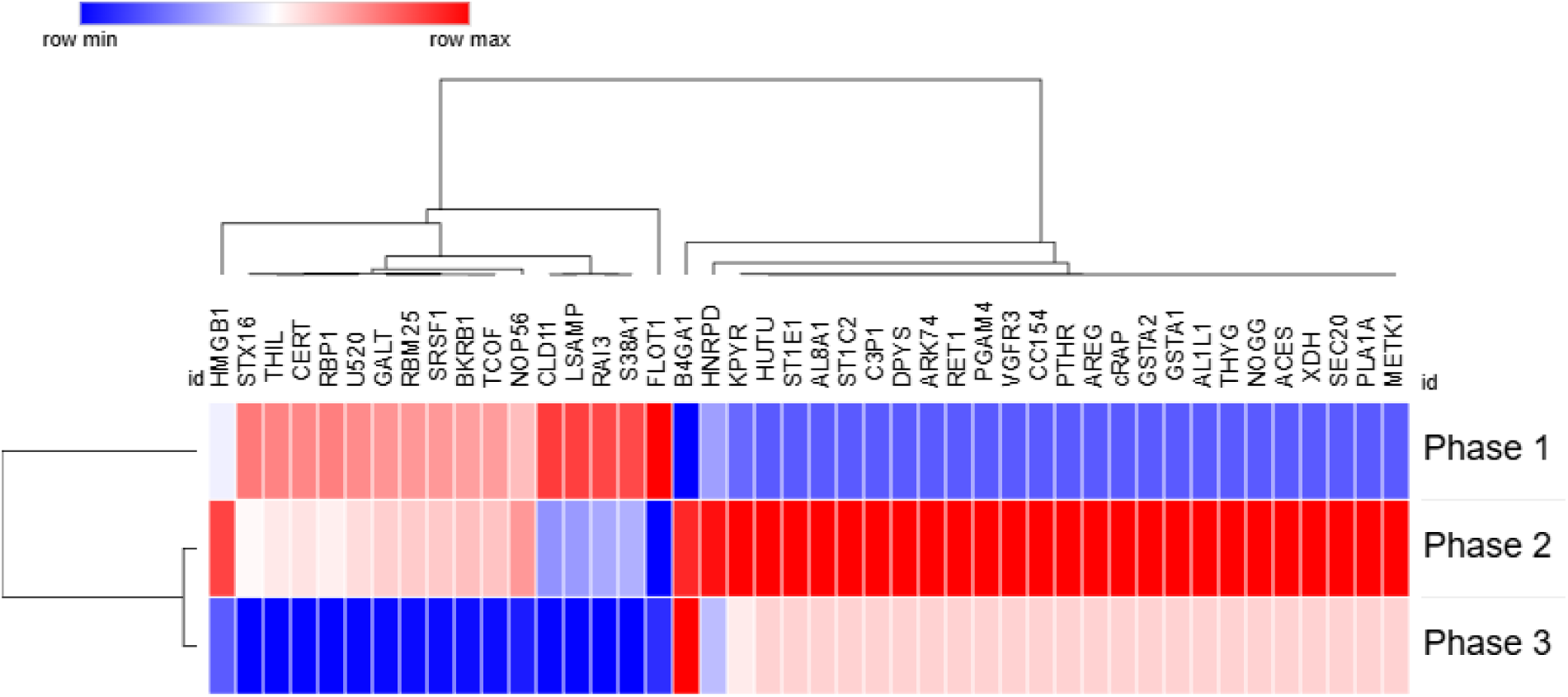
Transient phase-dependent remodeling of sEV protein cargo during prolonged HFB culture. Heatmap representation of exosomal proteins identified from Venn-intersection analyses showing non-monotonic, direction-switching regulation across culture phases. Rows correspond to individual proteins and columns to Phase 1, Phase 2, and Phase 3; protein abundance values are row-wise z-score normalized. Hierarchical clustering highlights distinct temporal expression patterns, including proteins that progressively decline from Phase 1 to Phase 3, predominantly associated with cell-cycle regulation, RNA processing, and cell–adhesion dynamics (e.g., HMGB1, SRSF1, RBM25, NOP56, STX16), as well as a second group displaying transient induction in Phase 2 followed by suppression in Phase 3, consistent with short-lived adaptive metabolic, stress-response, and paracrine signaling programs during senescence progression.

## Conclusions and future perspective

Prolonged culture within a hollow fiber bioreactor supports sustained expansion of human BMSCs while progressively reshaping their physiological state through cumulative microenvironmental and mechanical stress. Integrated cellular analyses including morphological assessment, senescence-associated β-galactosidase staining, confocal imaging, and flow cytometry revealed the emergence of an adaptive, early senescence–associated phenotype marked by cytoskeletal reorganization and reduced proliferative competence. Importantly, preservation of core mesenchymal surface marker expression indicated that lineage identity was largely maintained despite stress-induced aging. The sEVs closely mirrored these cellular transitions. NTA, protein quantification, and immunoblotting confirmed stable vesicle size distribution, marker expression, and production efficiency throughout extended culture, indicating preserved vesicle biogenesis. In contrast, quantitative proteomic profiling uncovered pronounced, stage-dependent remodeling of sEV cargo. Early culture phases were characterized by enrichment of proteins associated with metabolic and redox homeostasis, followed by increased representation of stress-response and cytoskeletal regulators, and culminating in the accumulation of extracellular matrix–associated and senescence-linked signaling components. This temporal progression delineates a continuum from cellular stress adaptation to stabilization of a senescence-associated secretory phenotype. Collectively, these findings indicate that long-term HFB culture promotes a controlled, stress-associated aging trajectory in BMSCs rather than acute functional deterioration. The close correspondence between cellular senescence phenotypes and sEV proteomic signatures establishes vesicle profiling as a sensitive, noninvasive, and integrative readout of stem cell aging in bioreactor systems. From a translational perspective, this convergence suggests that sEV analysis can capture key aspects of cellular aging while reducing reliance on parallel, labor-intensive cellular assays. More broadly, this work provides a mechanistic framework linking bioreactor-imposed stress, adaptive senescence, and dynamic extracellular vesicle remodeling, informing the rational design and monitoring of scalable stem cell culture platforms for regenerative and therapeutic applications.

## Author Contributions

Ali Dinari: Conceptualization, methodology, investigation, data analysis, visualization, writing original draft, review and editing. Armin M. Ebrahimi: Microscopy imaging and related data analysis. Bartosz Leszczyński: Data visualization and graphical analysis. Kamil Wawrowicz: Methodology at initial stages, data analysis, and related discussion. Masoud Rezaei: Methodology and imaging analysis. Maciej Stotwiński: Methodology. Zenon Rajfur: Microscopy imaging assessment. Ewa Stępień: Conceptualization, methodology, supervision, funding acquisition, writing, review and editing.

## Acknowledgements

We acknowledge Proteomics Core Facility of the Faculty of Biochemistry, Biophysics and Biotechnology and the Malopolska Centre of Biotechnology at the Jagiellonian University for mass spectrometry analysis. The purchase of Orbitrap Astral Mass Spectrometer has been supported by a grant from the Priority Research Area (POB BioS) under the Strategic Programme Excellence Initiative at Jagiellonian University.

